# Host community assembly modifies the relationship between host and parasite richness

**DOI:** 10.1101/857151

**Authors:** Fletcher W. Halliday, Robert W. Heckman, Peter A. Wilfahrt, Charles E. Mitchell

**Affiliations:** Department of Evolutionary Biology and Environmental Studies, University of Zurich, 8057, Zurich, CH; Department of Biology, University of North Carolina, Chapel Hill, NC 27599; Department of Integrative Biology, University of Texas at Austin, Austin, TX, USA 78712; Environment, Ecology and Energy Program, University of North Carolina, Chapel Hill, NC 27599; Department of Disturbance Ecology, University of Bayreuth, Bayreuth, Germany, 95447

**Author notes:** Correspondence: Winterthurerstrasse 190 8057 Zurich. **Author contributions:** FWH, RWH, and PAW designed and implemented the experiment. FWH analyzed the data and wrote the first draft. All authors contributed substantially to revising the manuscript.

**Keywords:** diversity-disease, parasite diversity, old fields, community assembly, phylogenetic diversity

## Abstract

Host and parasite richness are generally positively correlated, but the stability of this relationship during community assembly remains untested. The composition of host communities can alter parasite transmission, and the relationship between host and parasite richness is sensitive to parasite transmission. Thus, changes in composition during host community assembly could strengthen or weaken the relationship between host and parasite richness. Host community assembly, in turn, can be driven by many processes, including resource enrichment. To test the hypothesis that host community assembly can alter the relationship between host and parasite richness, we experimentally crossed host diversity and resource supply to hosts, then allowed communities to assemble. As previously shown, initial host diversity and resource supply determined the trajectory of host community assembly, altering post-assembly host species richness, richness-independent host phylogenetic diversity, and colonization by exotic host species. Throughout community assembly, host richness predicted parasite richness. As predicted, this effect was moderated by exotic abundance: communities dominated by exotic species exhibited a stronger positive relationship between post-assembly host and parasite richness. Ultimately, these results suggest that, by modulating parasite transmission, community assembly can modify the relationship between host and parasite richness, providing a novel mechanism to explain contingencies in this relationship.

## Introduction

Parasites are a major contributor to global biodiversity, yet parasite diversity remains relatively underexplored, a problem that has spurred recent research into the drivers of parasite diversity within host communities (Dobson *et al*. 2008; Kamiya *et al*. 2014; Johnson *et al*. 2016; McDevitt-Galles *et al*. 2018). Through this research within host communities, the positive relationship between host and parasite species richness has become one of the most consistently documented relationships in disease ecology (Hechinger & Lafferty 2005; Lafferty 2012; Kamiya *et al*. 2014; Johnson *et al*. 2016; Liu *et al*. 2016). Yet, whether this relationship is robust to changes in host community structure over time remains poorly understood, because few studies have quantified the relationship as host communities assemble. During community assembly, the structure of host communities shifts over time in response to biotic factors, such as species interactions, and abiotic factors, such as resource supply to hosts (HilleRisLambers *et al*. 2011; Harpole *et al*. 2016). These shifts in host community structure can alter parasite transmission (Johnson *et al*. 2013; Halliday *et al*. 2019), which could, in turn, alter the strength or direction of the relationship between host and parasite richness. Thus, the relationship between host and parasite species richness might depend on the manner in which host communities assemble (Johnson *et al*. 2016) (Fig 1).

**Fig. 1.**
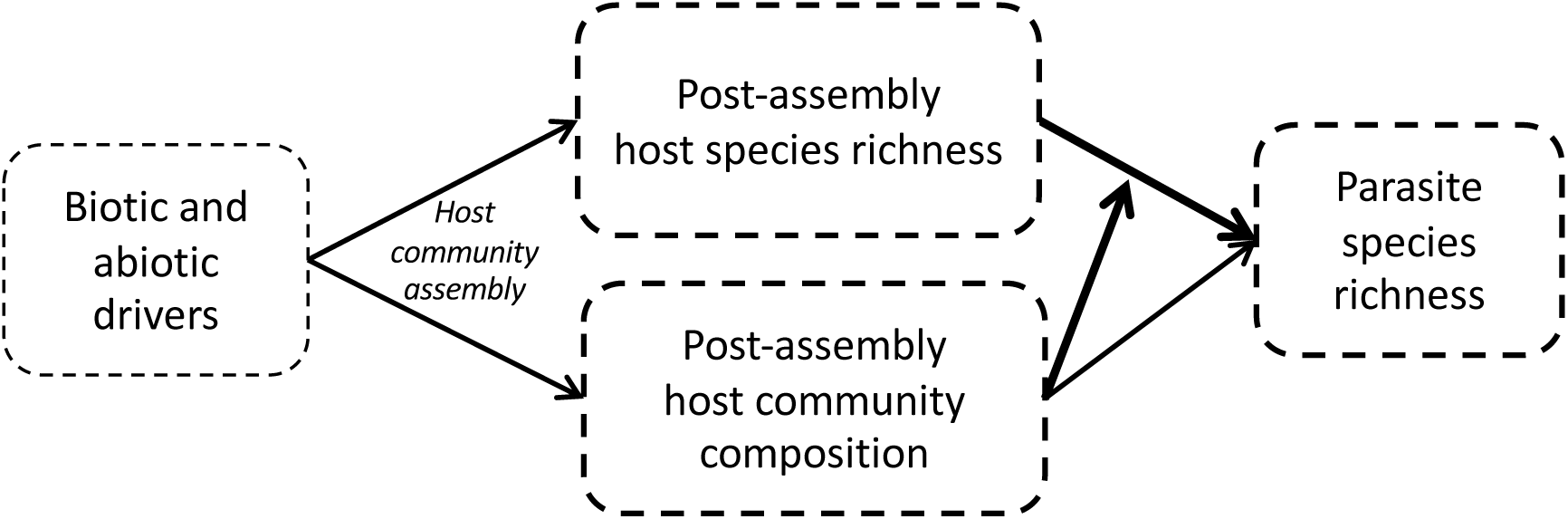
Conceptual metamodel of community assembly altering the relationship between host and parasite richness. During community assembly, biotic and abiotic drivers can determine the trajectory of host community composition, which may, in turn, moderate the relationship between host and parasite richness.

Within host communities, the “host-diversity-begets-parasite-diversity hypothesis” suggests that, because many parasites are specialized to infect a small number of host species, increases in host species diversity should increase parasite diversity (Hechinger & Lafferty 2005; Poulin 2014). Empirical support for this hypothesis is widespread. For example, a 2014 meta-analysis found a consistent positive relationship between host and parasite richness across 21 published studies (Kamiya *et al*. 2014). However, two recent studies highlight the possibility for important contingencies in the nature of this relationship. Wood et al (2018) observed that human activity decoupled the positive relationship between host and parasite richness among reef fishes, attributing the contingency in the relationship between host and parasite richness to a loss of parasite species with complex life-cycles. In another study, Johnson et al (2016) observed that within host communities, the positive relationship between host and parasite richness depended on the spatial scale of observation, attributing this contingency to colonization opportunities (e.g., the propagule-pressure hypothesis; Levine 2000), which weaken at larger spatial scales. Colonization opportunities are fundamental to metacommunity theory, suggesting that this contingency may be further understood by considering parasites in that theoretical framework.

Metacommunity theory predicts that, all else being equal, regional richness can be strongly influenced by dispersal and establishment of species among patches (Leibold *et al*. 2004; Holyoak *et al*. 2005; Logue *et al*. 2011). For parasites, transmission is the combination of dispersal and establishment, so multiple parasite species in a host population or community function as a metacommunity connected via transmission among host individuals (Kuris *et al*. 1980; Sousa 1994; Mihaljevic 2012; Borer *et al*. 2016; Mihaljevic *et al*. 2018). As such, metacommunity theory can be used to generate predictions about the relationship between host and parasite richness. When dispersal and establishment (i.e. parasite transmission) among patches (i.e., host individuals) is most limited, an “extinction vortex” can occur as rare species are lost from local patches faster than they can colonize new patches, leading to the eventual loss of those species in the entire metacommunity (i.e., host community) (Gilpin & Soule 1986). Greater extinction rates of parasites in host communities with low transmission could weaken the positive relationship between host and parasite richness. In a metacommunity context, reducing dispersal limitation by increasing habitat connectivity can alleviate this effect (Logue *et al*. 2011; Cornell & Harrison 2013). Similarly, when parasite metacommunities are transmission-limited, an increase in parasite transmission should strengthen the positive relationship between host and parasite richness (Poulin 2004). Thus, the magnitude of parasite transmission among hosts could explain variation in the relationship between host and parasite richness during host community assembly.

Host community assembly involves change over time in community characteristics that can alter parasite transmission, including host species richness, exotic host abundance, and host phylogenetic diversity (Box 1) (Halliday *et al*. 2019). These changes in host community characteristics may, in turn, be driven by a variety of biotic and abiotic conditions including initial host richness and resource supply to hosts (Fig. 1). Consequently, the impacts of initial biotic and abiotic conditions on host richness, parasite richness, and their interaction may shift as host communities assemble.

This study examined whether the relationship between host and parasite richness in a North Carolina old-field shifted during experimental host community assembly. Specifically, we constructed experimental, native plant communities at two initial host diversity levels and two levels of soil fertility, then measured how changes in post-assembly host richness, exotic host abundance, and host phylogenetic diversity (together describing host community assembly) influenced changes in parasite richness over three years. In previous analyses using a subset of our data, we found that initial host diversity and resource supply to hosts strongly influenced host community assembly, with consequences for exotic invasions, disease risk, and host community trait distributions (Heckman *et al*. 2017; Halliday *et al*. 2019; Wilfahrt *et al*. 2019). Here, we show that, despite strong shifts in the structure of host communities, host diversity begets parasite diversity during community assembly. Importantly however, the magnitude of the relationship between host and parasite richness depends on characteristics of the host community that are linked to parasite transmission.

## Methods

We performed this study in an old-field in Duke Forest Teaching and Research Laboratory (Orange County, NC, USA), dominated by perennial, herbaceous plants. To test whether initial host diversity and resource supply to hosts indirectly affect parasite richness via changes in host community assembly, we experimentally manipulated native plant (i.e., host) richness with multiple community compositions at each level of richness, and soil nutrient supply. This yielded a study that comprised 120 1m × 1m plots (5 replicate blocks × 2 resource supply levels × 2 host richness levels × 6 native community compositions). Because this study aimed to examine how plant and parasite community structure changes over time, we did not weed plots to maintain richness (Fargione & Tilman 2005; Heckman *et al*. 2017). Thus, the initial species richness treatments represent initial conditions and not the richness of a plot after July 2012. The full details of the experimental treatments can be found in Appendix A.

To establish initial host diversity, we assembled 12 planted communities at two richness levels from a pool of six species: six monocultures and six five-species polycultures where one species was excluded from each polyculture community. Plants were propagated from seed at the University of North Carolina at Chapel Hill, then seedlings were transplanted into the plots, spaced 10 cm apart.

We began resource supply treatments, hereafter referred to as the fertilization treatment, in July 2012. Each plot was either fertilized with 10 g m^-2^ each of N, P, and K or not fertilized. We applied slow-release forms of each nutrient each spring to alleviate nutrient limitation within experimental communities during the growing season (Borer *et al*. 2014).

### Quantification of host community structure

Each year, we visually quantified the percent cover of all plant species in each plot in September using a modified Daubenmire method (Borer *et al*. 2014). We evaluated changes in three components of host community structure to evaluate how experimental treatments influenced host community assembly: post-assembly plant species richness, exotic plant abundance, and the phylogenetic diversity of plant species. To quantify exotic plant abundance, we classified species as exotic or native to eastern North America using the USDA Plants Database (USDA & NRCS 2016), then assessed the relative abundance of exotic species (hereafter, exotic abundance) as the ratio of the absolute exotic cover to the total cover of all species within a plot. To quantify phylogenetic diversity, independent of species richness, a phylogeny of all non-tree species was constructed using ‘phyloGenerator’ (Pearse & Purvis 2013). Plant phylogenetic diversity was calculated using the ses.mpd function in R package Picante (Kembel *et al*. 2010). This function uses a null-modeling approach that measures the degree to which a plot is more or less phylogenetically diverse than random, given the number of host species, weighted by their relative abundance.

### Quantification of parasite richness

Following Lafferty and Kuris (2002), we define a parasite as any organism that spends at least one life history stage living in or on a single host individual, causing a fitness loss to the host. This definition includes all microbial parasites of plants and certain insect parasites of plants, such as galling and leaf-mining insects, which spend a larval life history stage parasitizing a single host leaf, but are transmitted by free-living adults that can seek out host plants for their offspring (Halliday *et al*. 2017a, 2019).

Parasites were surveyed in each plot annually for three growing seasons in September, which is the period of greatest parasite abundance in this system (Halliday *et al*. 2017b). In 2012, parasites were only surveyed on the six planted species (these species accounted for a median of 78% of total vegetative cover per plot), and were measured by haphazardly surveying five leaves on five individuals of each planted species in each plot.

In 2013, the composition of plots was characterized by a few common host species and many rare host species (Heckman *et al*. 2017). Because common host species contribute more to parasite abundance (Mordecai 2011; Heckman *et al*. 2016), parasites were measured by haphazardly surveying one individual of each of the six most abundant non-planted species across the experiment, as well as one individual of each of the six planted species that were surveyed in 2012 (together, the surveyed species accounted for a median of 78% of total vegetative cover per plot).

By 2014, the composition of host communities had shifted considerably from the originally planted compositions (Heckman *et al*. 2017; Halliday *et al*. 2019; Wilfahrt *et al*. 2019). To estimate parasite richness in the assembled community, we maximized the number of host species surveyed in a plot and across the experiment, by haphazardly surveying parasites on five leaves on one individual of the most abundant host species, and then the next most abundant species, iterating until the sampled species’ summed cover accounted for at least 80% of the plot’s total plant cover. We surveyed one additional individual of all six planted host species in each plot, regardless of cover. While this sampling method reduces replication at the scale of host individuals within plots, it samples across leaf ages and matches similar approaches for measuring community-wide responses at the plot scale (e.g., Pérez-Harguindeguy *et al*. 2013). Visual surveys were conducted following Halliday et al (2017a, 2019). Briefly, parasites were categorized into morphospecies using morphological and genetic characteristics (Table S1).

The number of host species, and thus individuals, surveyed varied among plots (min = 2, median = 5, mean = 7.9, max = 25 host individuals). We therefore performed “site-based” rarefaction on the count of parasite morphospecies (Gotelli & Colwell 2001). In each plot, we randomly sampled up to five host individuals, and counted the number of parasite morphospecies in that subsample. We permuted this 999 times and took the average rarefied parasite richness for each plot across those 999 permutations. Although insect and microbial parasites may respond differently to host richness and fertilization (Halliday *et al*. 2017a), the post-assembly data on insects were not sufficient to test for differences from microbes. Therefore, rarefied parasite richness was calculated across all parasites, including insects and microbes.

### Data analysis

#### Longitudinal model of parasite richness

We first tested whether the relationship between host and parasite richness would change over time as a function of initial host diversity, fertilization, and their interaction by constructing a longitudinal linear mixed model using the lme function in the nlme package (Pinheiro *et al*. 2016). In order to meet assumptions of homoscedasticity, we added an identity variance structure (varIdent function) by host diversity treatment (Zuur *et al*. 2009; Pinheiro *et al*. 2016). Each model included the fertilization treatment, the initial host diversity treatment, post-assembly host species richness, year of observation, and all interactions between these four variables as fixed effects, plus block as a non-interacting fixed covariate. To account for temporal autocorrelation, we included an AR 1 autocorrelation structure at the plot level in each model (Zuur *et al*. 2009). We included planted composition as a random effect to ascribe differences to initial host diversity only when differences in a response within a richness level (i.e., polycultures or monocultures) were smaller than differences between richness levels (Schmid *et al*. 2002). This analysis tests the effect of the initial host diversity treatment after accounting for variation in host composition. We used the pairs function in the lsmeans package (Lenth 2016) to test whether treatment means were different in a given year, and used the lstrends function to test whether the slope of the relationship between post-assembly host and parasite richness differed among treatments and years. However, we caution the comparison of treatment means among years, as sampling methodology for parasites varied between years, as detailed above. In order to facilitate comparisons among responses and clarify relationships among predictors, we reduced the model by removing non-significant interactions (following Crawley 2007; Zuur *et al*. 2009).

#### Structural equation model including community assembly

The longitudinal model of parasite richness tested whether the relationship between post-assembly host and parasite richness changed over time as a function of experimental treatments. To explicitly test the hypothesis that the relationship between post-assembly host and parasite richness is altered by shifts in host community structure during community assembly, we performed confirmatory path analysis using the lavaan package (Rosseel 2010). Specifically, we fit a structural equation model (SEM) that included the treatment effects and their interaction on three endogenous mediators (post-assembly host species richness, exotic host abundance, and host phylogenetic diversity) measuring the outcome of community assembly (following Halliday *et al*. 2019), as well as the effects of those three mediators on parasite richness (Box Fig. 1). We tested the hypothesis that shifts in host community structure altered the relationship between post-assembly host and parasite richness by fitting a second-stage moderated mediation (Hayes 2015) including pairwise interactions between exotic host abundance and post-assembly host richness (Box Fig 1 path m) and between host phylogenetic diversity and post-assembly host richness (Box Fig 1 path n). The first half of this model (Box Fig 1 paths a-i) uses the same data and is structurally identical to the paths from the 2014 group in the multigroup model presented in Halliday et al (2019).

Experimental block was treated as a stratified independent grouping variable using the lavaan.survey package (Oberski 2014). In order to meet assumptions of homoscedasticity and multinormality, we logit transformed exotic abundance. All endogenous variables were mean centered, following transformation, to improve interpretability and to eliminate non-essential collinearity (Toothaker *et al*. 1991). In order to facilitate comparisons among responses and clarify relationships among predictors, we reduced the model by removing non-significant interactions (following Crawley 2007; Zuur *et al*. 2009), limiting model reduction to exogenous (i.e., treatment) variables only.

## Results

### Longitudinal model of parasite richness

We first tested the hypothesis that the relationship between host and parasite richness would change over time as a function of initial biotic and abiotic conditions by constructing a longitudinal mixed model of parasite richness as a function of post-assembly host species richness and its interaction with initial host diversity and fertilization. Consistent with our hypothesis, there was a positive relationship between host and parasite richness, which changed over time as a function of initial host diversity (Appendix B; Table S1). However, fertilization did not affect the relationship between host and parasite richness over time (Table S2), and so these interactions were removed from the model, yielding a reduced model (Table S3).

In the reduced model, initial host diversity altered parasite richness, and this effect varied over time (year p=0.001; initial host diversity × year p<0.0001; Table S3; Fig 2). In 2012, parasite richness was more than twice as high in polycultures as in monocultures (p<0.001), providing strong evidence for the hypothesis that host diversity begets parasite diversity, though that effect was reduced by 40% in experimentally fertilized communities (fertilization × initial host diversity p=0.003). The positive effect of initial host diversity on parasite richness weakened in 2013 (p=0.014), and by 2014, as the communities were further colonized by non-planted host species, the positive effect from previous years became a negative effect, though only in fertilized communities: parasite richness was 17% lower in fertilized polycultures than in fertilized monocultures.

**Fig. 2.**
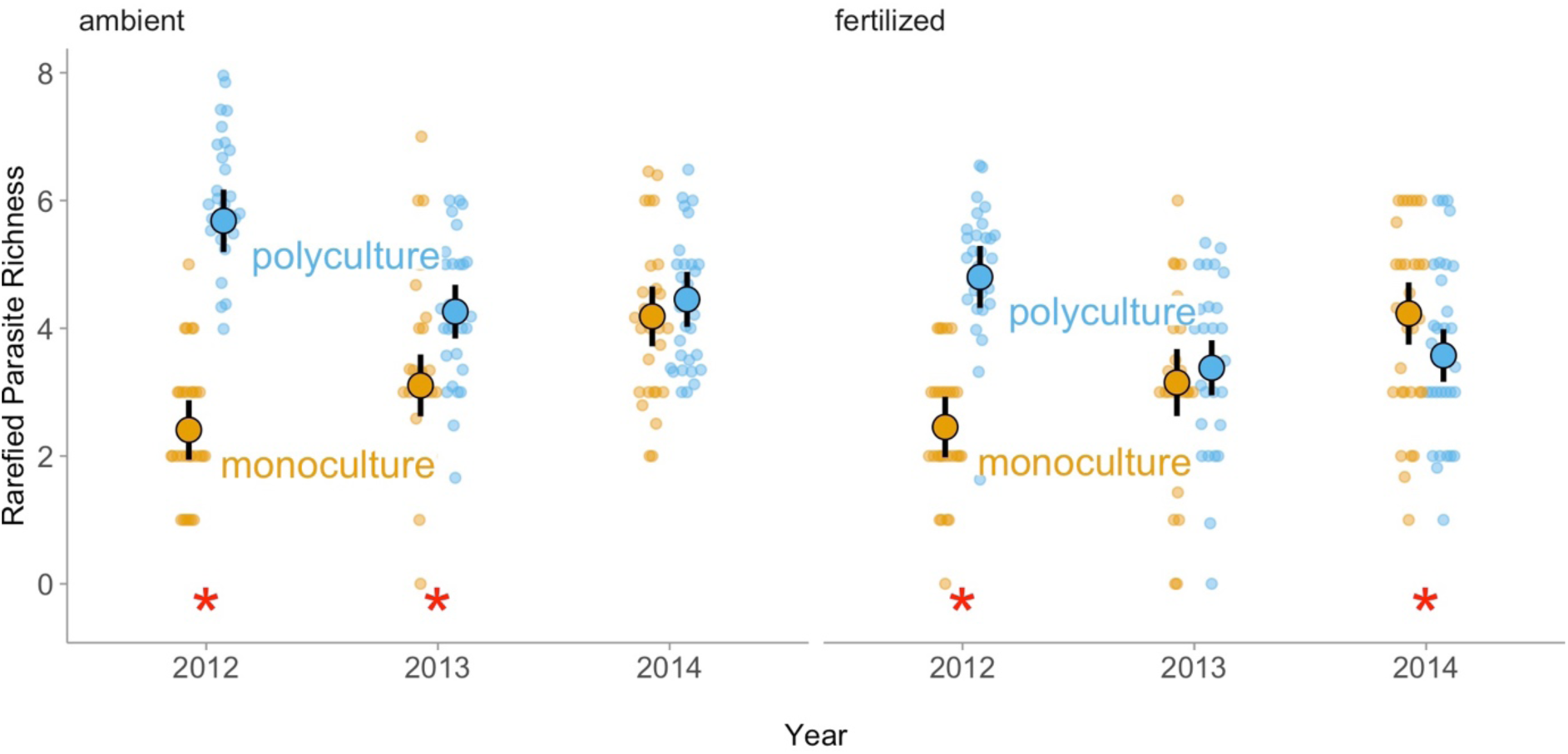
Longitudinal mixed model results showing the model-estimated effects of initial host richness (monocultures orange; polycultures blue) and resource supply (unfertilized communities on the left; fertilized communities on the right) on rarefied parasite richness across three years. Large points represent the estimated treatment mean, error bars are 95% confidence intervals, and small points show the raw data. The y-axis shows rarefied parasite richness. Asterisks show significant effects (p < 0.05). Initial host richness increased parasite richness but that effect weakened as host communities assembled, and ultimately reversed in fertilized host communities.

In the reduced model, the positive relationship between host and parasite richness depended on initial host diversity (initial host diversity × post-assembly host richness p=0.04), and this effect of initial host diversity changed over time (initial host diversity × post-assembly host richness × year p=0.01; Fig 3; Fig S1; Table S3). In 2012, there was a significantly positive relationship between post-assembly host and parasite richness in polycultures (p=0.03). This effect became non-significant as host communities assembled over time (p_2013_=0.42; p_2014_=0.37). In contrast, there was no significant positive relationship between post-assembly host and parasite richness in monocultures in 2012 or 2013 (p_2012_=0.20; p_2013_=0.87). However, the relationship strengthened over time, such that by 2014, there was a significant positive relationship between post-assembly host and parasite richness in monocultures (p=0.004). This change over time in the relationship between post-assembly host and parasite richness as a function of initial host diversity suggests that initial host diversity altered the trajectory of host community assembly over time, thereby altering the relationship between post-assembly host and parasite richness. The nature of this altered trajectory is explored in the structural equation model.

**Fig. 3.**
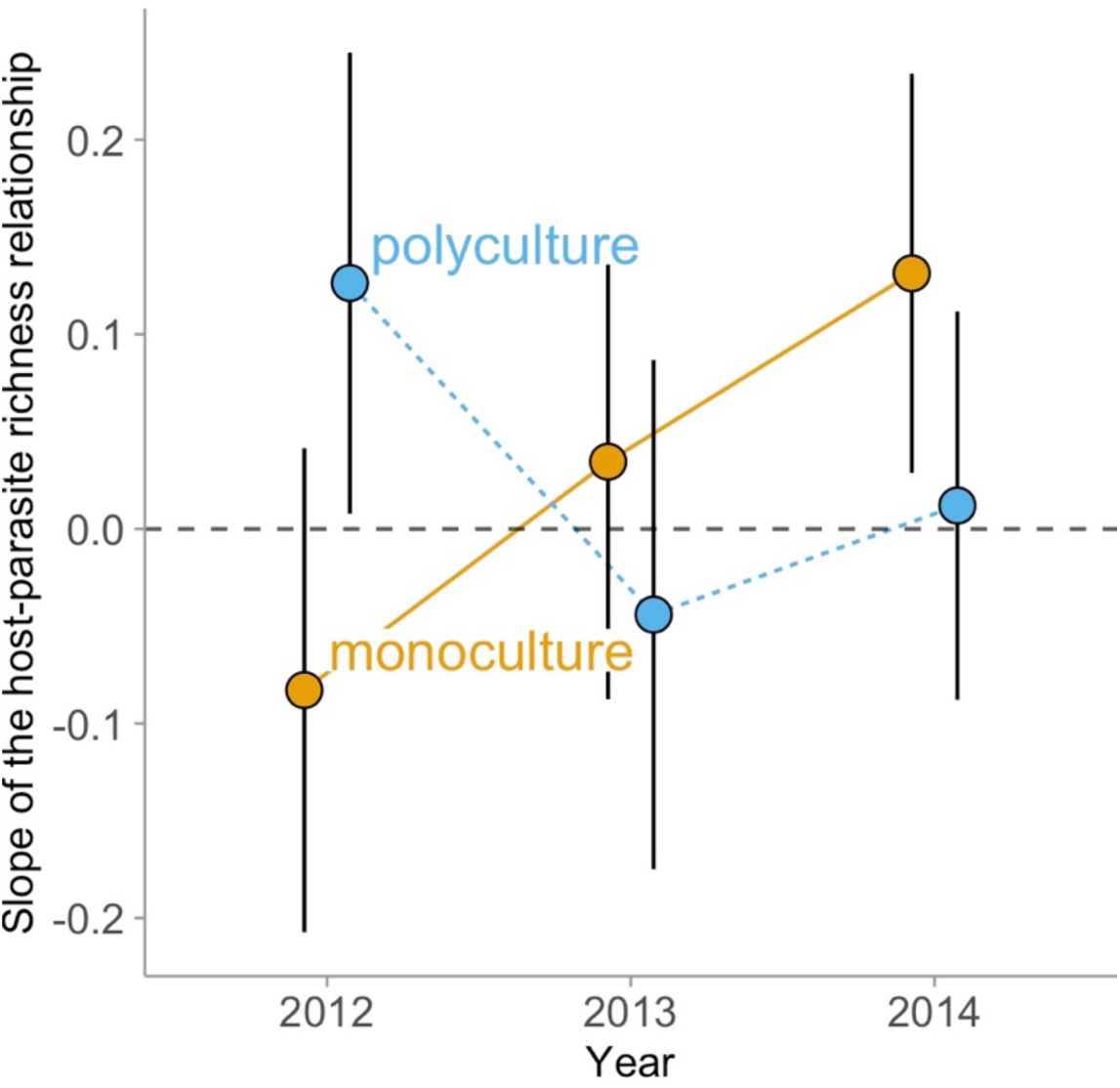
Longitudinal mixed model results showing the model-estimated effect of post-assembly host richness on rarefied parasite richness as a function of initial host diversity (monocultures orange; polycultures blue) across three years. The y-axis shows the slope of the relationship between post-assembly host richness and parasite richness. Points represent the estimated effect of the experimental treatment on that slope (i.e., the interactive effect of post-assembly host richness and initial host diversity on parasite richness). Error bars are 95% confidence intervals. The positive relationship between host and parasite richness that was observed in polycultures in 2012 weakened over time, while the relationship between host and parasite richness in monocultures strengthened over time, resulting in a positive relationship in 2014.

### Structural equation model including community assembly

The longitudinal mixed model of parasite richness indicated that the relationship between post-assembly host richness and parasite richness changed over time as a function of initial host diversity. That change may have been driven by shifts in host community structure (i.e. community assembly). We therefore next tested the hypothesis that initial host diversity and fertilization indirectly influenced post-assembly parasite richness via their impacts on post-assembly host richness, exotic abundance, phylogenetic diversity, and their interactions, using a structural equation model (Box 1). The data were well fit by this model (Robust χ^2^ p=0.288, RMSEA p=0.618, SRMR=0.075), though initial host diversity and fertilization did not interactively influence any of the response variables (p>0.05; Table S4). Therefore, we removed these interactions, yielding a reduced model (Fig 4, Table S5).

**Fig. 4.**
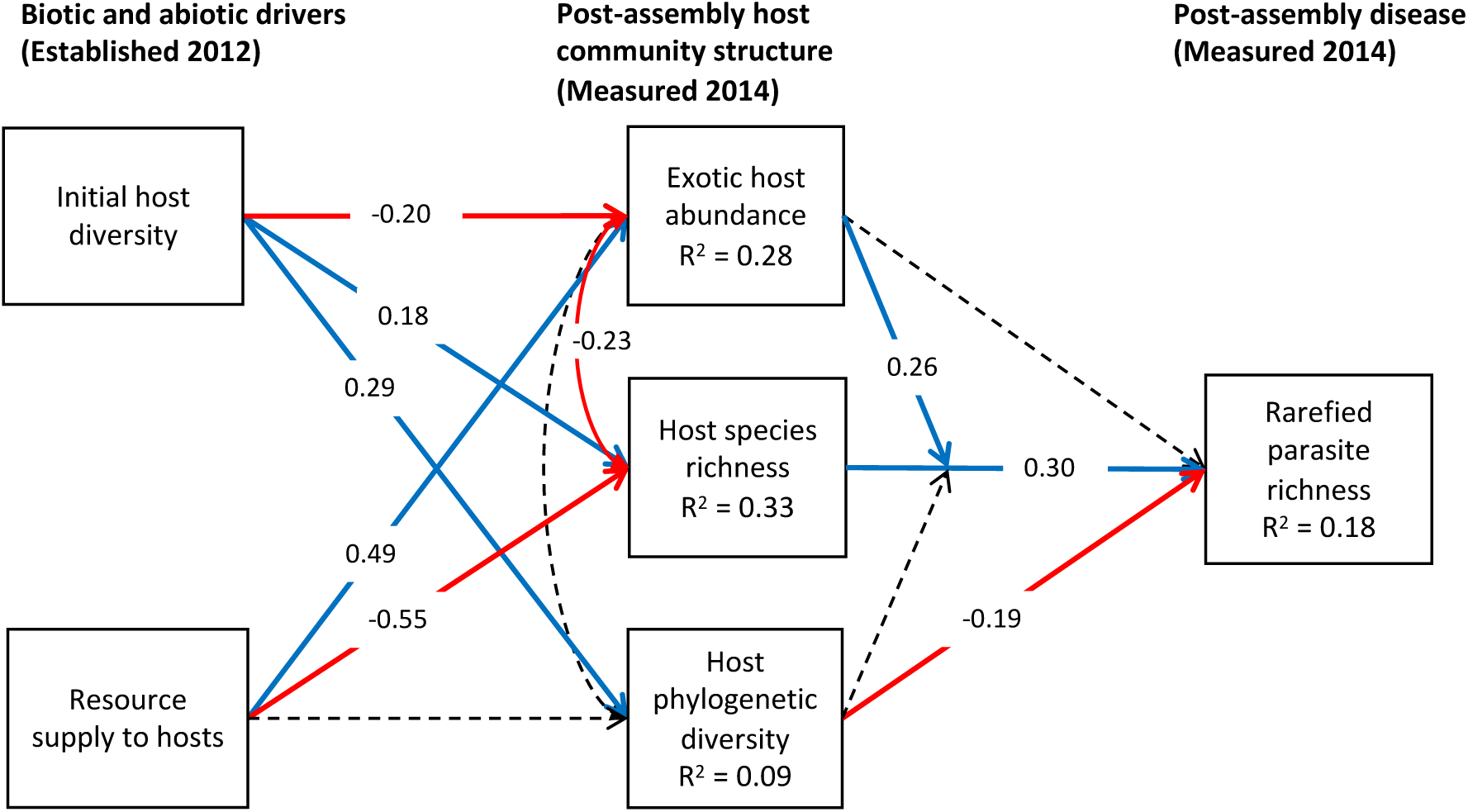
Structural equation model results for the final (reduced) model. Red lines are negative effects (p < 0.05), blue lines are positive effects (p < 0.05), and dashed lines are non-significant (p > 0.05). All coefficients are standardized. Correlations between errors are denoted with double-headed arrows. Higher post-assembly host richness increased parasite richness, and that effect was stronger in communities that, as a result of community assembly, became more heavily dominated by exotic host species.

To model host community assembly, the structural equation model included structurally identical paths, and the same data as the 2014 group from the model presented in Halliday et al (2019). As shown in that model and in other results from this study system, initial biotic and abiotic conditions determined the trajectory of host community assembly, with host communities becoming increasingly divergent over time depending on initial biotic and abiotic conditions (Heckman *et al*. 2017; Halliday *et al*. 2019; Wilfahrt *et al*. 2019). Specifically, increasing initial host diversity increased post-assembly host richness, reduced exotic host abundance, and increased richness-independent phylogenetic diversity of host species. Fertilization reduced post-assembly host richness and increased exotic host abundance, but had no significant effect on phylogenetic diversity of host species (Halliday *et al*. 2019). Having re-analyzed this previously published first stage of the structural equation model to quantify the effects of experimental treatments on host community assembly, we then evaluated a novel second stage of the model (Box 1 Fig. 1) to test the hypothesis that shifts in host community structure drive and moderate the effect of post-assembly host richness on parasite richness.

As predicted, post-assembly host richness increased parasite richness (p=0.001), supporting the hypothesis that host diversity begets parasite diversity (Fig 4). This effect was moderated by shifting exotic abundance during community assembly: communities that became heavily dominated by exotic species also exhibited the strongest positive relationship between post-assembly host and parasite richness (p<0.0001; Fig 5; Fig S2), even though exotic abundance did not directly influence parasite richness (p = 0.16). Parasite richness was also lower in communities that became more phylogenetically overdispersed (p=0.026), consistent with previous observations that more phylogenetically overdispersed communities may exhibit lower parasite transmission. However, phylogenetic diversity did not alter the relationship between post-assembly host and parasite richness (p=0.40). Together these results indicate that, while the relationship between host and parasite richness is consistent, the magnitude of this relationship depends on community assembly. Consequently, these results support the hypothesis that host diversity begets parasite diversity, but reveal important contingencies in how this relationship manifests over time.

**Fig. 5.**
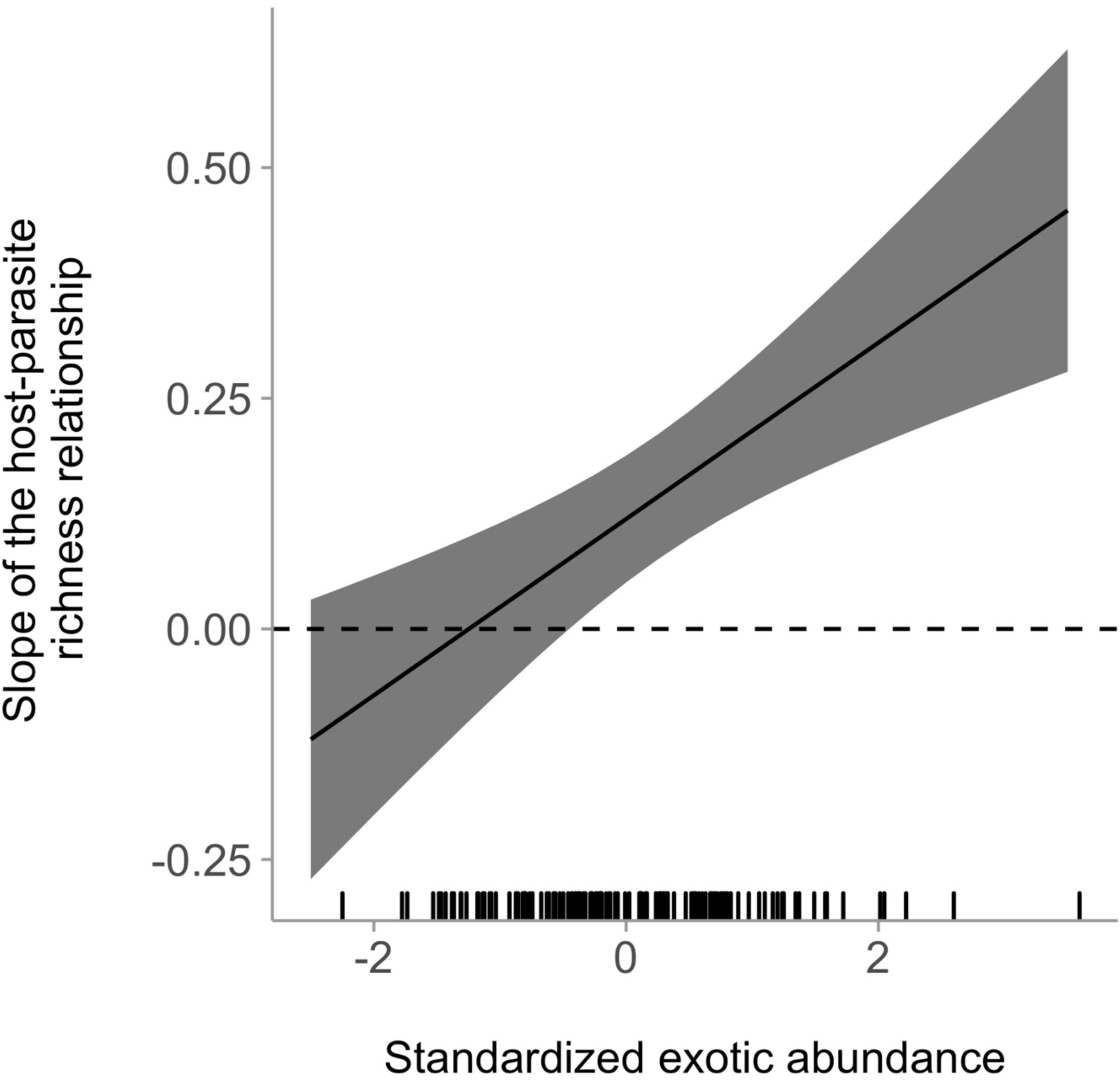
Effect of post-assembly host richness on parasite richness as a function of exotic host abundance. Model estimated effects of standardized and centered exotic abundance on the slope of the relationship between post-assembly host richness and parasite richness (i.e., the interactive effect of post-assembly host richness and exotic host abundance on parasite richness). The rug along the x-axis shows the distribution of the standardized and centered empirical data. Communities that fall 0.5 or more standard deviations below the mean of exotic host abundance show no relationship between host and parasite richness. Above that level, the relationship between host and parasite richness becomes increasingly positive with increasing exotic host abundance. Consequently, communities that became most dominated by exotic species also exhibited the strongest positive relationship between host and parasite richness.

## Discussion

In this study, the relationship between host and parasite richness changed over time as the host community assembled. Specifically, higher planted host diversity led to an initial increase in parasite richness that weakened as the host community assembled, and the ultimate relationship between post-assembly host and parasite richness was contingent on how host community structure changed during community assembly. This represents, to our knowledge, the first experimental evidence that the relationship between host and parasite richness can change over time in response to shifting host community structure during host community assembly. This result supports two recent studies that found that the relationship between host and parasite richness was contingent on characteristics of host and parasite communities (Johnson *et al*. 2016; Wood *et al*. 2018). Furthermore, host community assembly is linked to parasite transmission (e.g., Joseph *et al*. 2013; Johnson *et al*. 2015; Halliday *et al*. 2019), and our results suggest that contingencies in the relationship between host and parasite richness during host community assembly can emerge as a consequence of changing parasite transmission.

Our hypothesis that changes in parasite transmission during host community assembly would modify the relationship between host and parasite richness was grounded in metacommunity theory. According to metacommunity theory, the richness of species in a metacommunity often depends on habitat heterogeneity, but this effect can be highly sensitive to reduced dispersal and establishment among patches (Leibold *et al*. 2004). A host community is a parasite metacommunity, with the richness of host species in the community representing heterogeneity of parasite habitat (Johnson *et al*. 2016), and parasite transmission representing dispersal and establishment of parasites among host individuals within that metacommunity (Borer *et al*. 2016). Thus, parasite richness should increase with increasing host richness, but this effect may be altered by parasite transmission. Parasite transmission, in turn, can be altered by shifts in host community structure that take place during community assembly (Johnson *et al*. 2013; Halliday *et al*. 2019). Exotic hosts contribute the most to disease in this system (Halliday *et al*. 2019; Heckman *et al*. 2019), and consequently, the abundance of exotic hosts modified the relationship between host and parasite richness, supporting our hypothesis.

The relationship between post-assembly host and parasite richness was contingent on the final abundance of exotic host species that resulted from community assembly. Many of the exotic hosts that dominate Southeastern US old fields were introduced by humans from fertilized pastures (Fridley 2008), benefit from experimental fertilization (Heckman *et al*. 2016, 2017), and are sensitive to initial host diversity. Previous analyses indicate that exotic hosts also contributed most to parasite abundance in communities that they dominated, suggesting that exotic host species may contribute disproportionately to parasite transmission (Halliday *et al*. 2019). Consequently, we suggest that increasing exotic abundance may have strengthened the relationship between host and parasite richness by alleviating dispersal limitation of parasites in communities dominated by exotic species. This hypothesis, that exotic host species contribute disproportionately to parasite transmission, relies on the assumption that native and exotic hosts can share at least some of the same parasite species. Nearly half (6/13) of the parasites infecting exotic host species were also observed infecting native host species, providing some support for this assumption (Table S1).

The effect of exotic abundance on the relationship between host and parasite richness is consistent with predictions grounded in metacommunity theory (Leibold *et al*. 2004; Holyoak *et al*. 2005), but could this pattern be driven by the design of this study? In this system, high exotic abundance typically results from increasing abundance of a few dominant species (Heckman *et al*. 2017), suggesting strong competitive asymmetry. Consequently, these dominant species are capable of reducing the abundance and richness of other species (MacDougall *et al*. 2009), which, in addition to increasing parasite transmission, could interfere with how parasite richness was estimated. If this were occurring here, we would have under-sampled parasite diversity in heavily invaded host communities. But we found no difference between the percent of host species surveyed in communities that were minimally invaded (at least one standard deviation below mean exotic abundance; mean ± SD: 77 ± 10%) and communities that were heavily invaded (at least one standard deviation above mean exotic abundance; mean ± SD: 83 ± 11%). This indicates that under-sampling of parasite diversity in heavily invaded host communities was unlikely to contribute to our observed results. In other systems, though, high exotic abundance can arise from increasing the abundance of many sub-dominant species. These exotic species typically coexist with native species by occupying distinct niches (MacDougall *et al*. 2009), which should have less negative effects on the abundance and richness of other species. This may lead to increased diversity in invaded host communities (Fridley *et al*. 2007), which could reduce parasite transmission. Future studies could resolve these potential issues by experimentally manipulating exotic abundance (or other drivers of parasite transmission) and host richness simultaneously (Young *et al*. 2017). Nevertheless, these results highlight the value of our dynamic community assembly approach for generating new hypotheses that could be tested through direct manipulations of static host communities.

Increasing host phylogenetic diversity reduced parasite richness, consistent with previous observations that more phylogenetically overdispersed communities may exhibit lower parasite transmission (e.g., Parker *et al*. 2015; Halliday *et al*. 2019). These results are in contrast to predictions grounded in the phylogenetic signal in parasites’ host range (Box 1), which suggest that, because more distantly related hosts are less likely to share pathogens than closely related host species (Gilbert & Webb 2007), communities with higher phylogenetic diversity may support more parasite species (e.g. via habitat heterogeneity; Johnson *et al*. 2016). Experiments that explicitly cross host taxonomic and phylogenetic diversity (e.g., Cadotte 2013) would help disentangle the conditions under which taxonomic and phylogenetic diversity can have opposite, but non-interacting results.

Our hypothesis that increasing transmission during host community assembly would alleviate dispersal limitation for parasites relies on the assumption that parasite communities would be dispersal limited. This assumption appears to hold for some parasites of plants (Mitchell *et al*. 2002; Laine 2005; Tack *et al*. 2014; Halliday *et al*. 2017a) and animals (Sousa 1994; Esch *et al*. 2001; Byers *et al*. 2008; Mihaljevic *et al*. 2018). However, even in systems where parasite dispersal does not appear to be particularly limiting (e.g., Richgels *et al*. 2013; Dallas & Presley 2014; Ekholm *et al*. 2017), increasing transmission could still alter the relationship between host and parasite richness. In a metacommunity, when species richness is not dispersal-limited, mass effects (Logue *et al*. 2011; Cornell & Harrison 2013) can reduce regional richness by overwhelming species sorting mechanisms that occur within habitat patches. Similarly, when parasite distributions are not transmission-limited, an increase in parasite transmission could weaken the relationship between host and parasite richness (e.g., via mass effects), particularly if host generalists replace more specialized parasites (Johnson *et al*. 2016). Thus, across a wide gradient in parasite transmission rate, we hypothesize the effect of transmission rate on the relationship between host and parasite richness to be nonlinear: a positive/strengthening effect at low transmission rate, and a negative/weakening effect at high transmission rate.

While our results demonstrate that the relationship between host and parasite richness can be moderated by shifting host community structure, they are nonetheless consistent with the well-established “host-diversity-begets-parasite-diversity” hypothesis (Hechinger & Lafferty 2005; Kamiya *et al*. 2014) . Consistent with previous experiments in other systems, host richness most strongly predicted parasite richness immediately following establishment of experimental host communities (Rottstock *et al*. 2014; Liu *et al*. 2016). This effect weakened as non-planted hosts colonized experimental communities, but a significant positive effect of post-assembly host and parasite richness remained even after two years of community assembly. This latter result is consistent with the “host-diversity-begets-parasite-diversity” hypothesis (Hechinger & Lafferty 2005), even after accounting for the moderating effect of community assembly.

Together, these results demonstrate that the relationship between host and parasite richness can be contingent on characteristics of host community structure that shift during host community assembly. Thus by leveraging host community assembly, this study adds a novel mechanism to a growing body of literature revealing key contingencies in this relationship (Johnson *et al*. 2016; Wood *et al*. 2018). We suggest that contingencies in the relationship between host and parasite richness may occur because characteristics of host and parasite communities that shift during community assembly alter parasite transmission. More specifically, parasite transmission may be altered simultaneously by multiple components of host community structure that during community assembly shift in concert. For example, experimental fertilization both increased colonization by exotic host species and decreased host species richness (Heckman *et al*. 2017; Halliday *et al*. 2019). This covariance, in turn, led to an indirect effect on the relationship between host and parasite richness: communities that were more heavily invaded exhibited a stronger positive relationship between host and parasite richness. Consequently, fertilization indirectly strengthened the relationship between host and parasite richness. This study represents an important step forward by providing an ecological mechanism to explain contingencies in one of the most consistently reported patterns in disease ecology – the positive relationship between host and parasite diversity.

## Acknowledgments

J. Bruno, J. Wright, S. Halliday, and members of the Mitchell Lab provided helpful suggestions for the design of this experiment and assistance with plant propagation, fieldwork, and data analysis. We are also grateful for insightful suggestions from A-L. Laine and members of the Laine Lab. FWH, RWH, and PAW were supported by UNC’s Alma Holland Beers Scholarship and WC Coker Fellowship. RWH and FWH were supported by the UNC Graduate School Dissertation Completion Fellowship. FWH was supported by the NSF Graduate Research Fellowship. This work was supported by the NSF-USDA joint program in Ecology and Evolution of Infectious Diseases (NSF grant DEB-1015909 and USDA-NIFA AFRI grant 2016-67013-25762).

## Box 1.

### Biotic and abiotic drivers of host community assembly can indirectly alter the host-parasite richness relationship

Host community assembly involves change over time in community characteristics in response to a variety of biotic and abiotic conditions. Here, we consider three characteristics of host communities that change over time during community assembly and can alter parasite transmission: host species richness, exotic host abundance, and host phylogenetic diversity. We consider these changes in host community structure in response to two potential drivers: initial host diversity and resource supply to hosts.

Increased resource supply often reduces host species richness by decreasing the number of limiting resources that species compete for (Box Fig. 1, path h; Harpole *et al*. 2016), increases the abundance of exotic species by favoring species adapted to resource-rich environments (Box Fig. 1, path g; Huenneke *et al*. 1990; Heckman *et al*. 2017), and reduces phylogenetic diversity by favoring clades with specific resource uptake and allocation strategies (Box Fig. 1, path i; Mayfield & Levine 2010). Furthermore, communities that assemble from higher initial diversity may experience a legacy effect, with these host communities maintaining higher richness during community assembly (Box Fig. 1, path b; Mouquet *et al*. 2003), being more likely to resist invasion by exotic species at small and intermediate spatial scales (Box Fig. 1, path a; Levine & D’Antonio 1999; Fargione & Tilman 2005), and promoting colonization by species from different clades with low niche overlap, resulting in increased phylogenetic overdispersion (Box Fig. 1, path c; Mayfield & Levine 2010; Pavoine & Bonsall 2011). By altering parasite transmission, these concurrent shifts in host community structure might alter the relationship between host and parasite richness as host communities assemble.

Shifts in exotic host abundance during community assembly might alter the relationship between host and parasite richness. Parasite richness often increases with host richness, because higher host richness represents a more diverse pool of resources for parasites (Box Fig. 1, path k; Kamiya *et al*. 2014). However, exotic species often initially escape the parasites that infected them in their native range (Mitchell & Power 2003; Mitchell *et al*. 2010; Heger & Jeschke 2014), potentially leading to lower parasite richness (Box Fig. 1, path j) and weakening the relationship between host and parasite richness (Box Fig. 1, path m) by reducing transmission in exotic-dominated communities. Alternatively, introduced hosts can also acquire infections from closely related native hosts (Parker *et al*. 2015) or via repeated introductions over time (Mitchell *et al*. 2010; Stricker *et al*. 2016). Successful exotic species are often more competent hosts for the parasites that can infect them (Han *et al*. 2015; Young *et al*. 2017), which could increase parasite transmission in exotic-dominated communities (e.g., Halliday *et al*. 2019). Thus shifts in exotic host abundance could strengthen or weaken the relationship between host and parasite richness, depending on characteristics of the host and parasite communities (Box Fig 1, path m).

Shifts in host phylogenetic diversity during community assembly might alter the relationship between host and parasite richness. More distantly related hosts are less likely to share pathogens (Gilbert & Webb 2007). Consequently, communities with higher host phylogenetic diversity, independent of species richness, may support more parasite species (Box Fig. 1, path l), potentially strengthening the relationship between host and parasite richness in phylogenetically overdispersed host communities (Box Fig. 1, path n). However, as host phylogenetic overdispersion increases, parasite transmission is expected to decline via a phylogenetic dilution effect (Parker *et al*. 2015; Liu *et al*. 2016; Halliday *et al*. 2019), which could reduce parasite species richness and weaken the relationship between host and parasite richness in phylogenetically overdispersed host communities (Box Fig. 1, paths n, l). Thus, drivers of community assembly that favor higher phylogenetic overdispersion could either increase or reduce parasite richness, while simultaneously strengthening or weakening the relationship between host and parasite richness.

**Box Fig 1.**
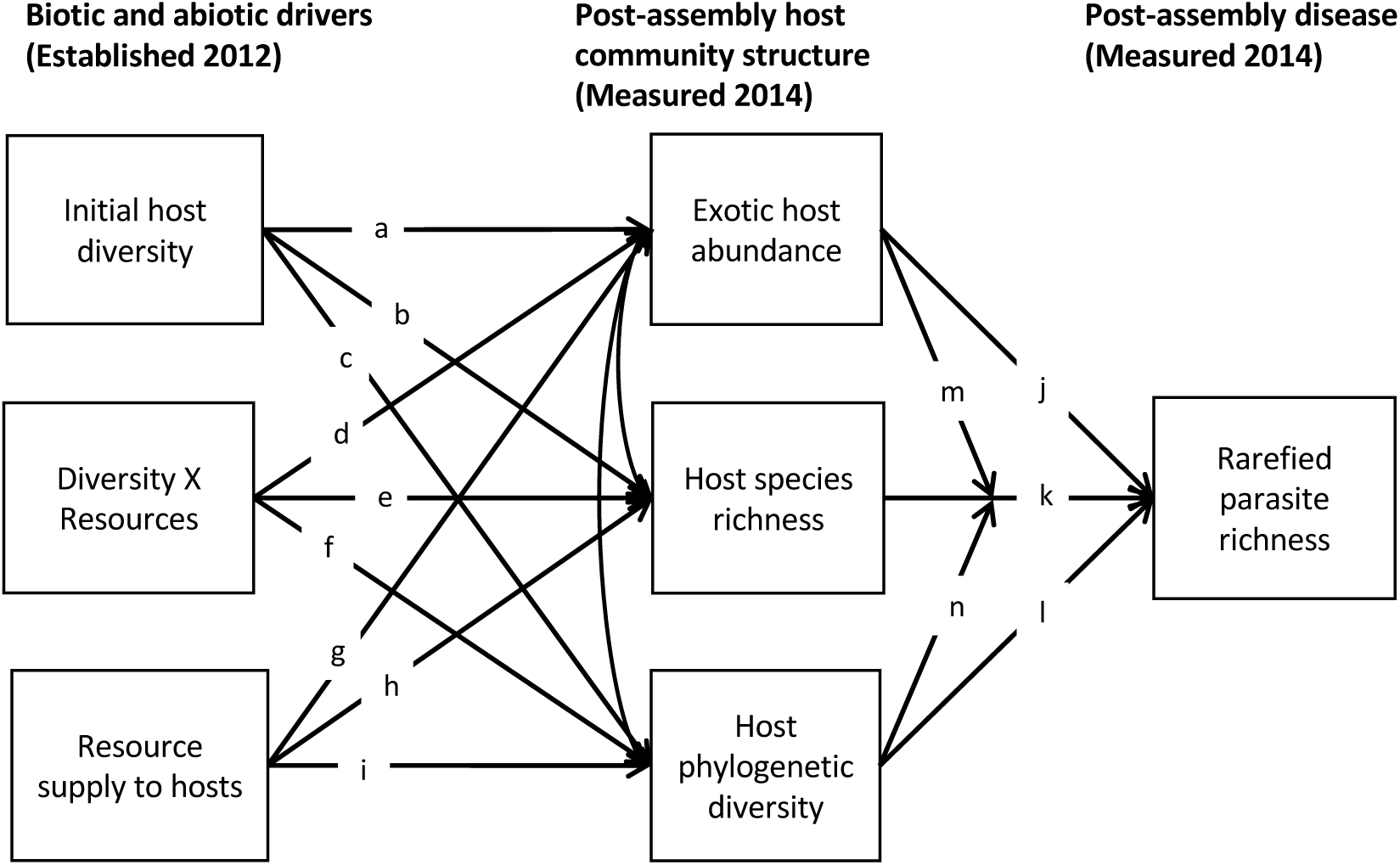
Measurement model representing the hypothesized effects of initial host diversity and resource supply on parasite richness, mediated by post-assembly host community structure. Paths are labeled a-n for reference in the text. Paths a - i represent the effects of experimental treatments on community assembly (the model’s first stage). Paths j - n (the model’s second stage) test the hypothesis that shifts in host community structure altered the relationship between post-assembly host and parasite richness. Specifically, paths j - l represent direct effects of the final plant community in 2014 on parasite richness. Paths m - n represent the moderating effect of shifting community structure on the relationship between host and parasite richness.

