## Appendix for "Host community assembly modifies the relationship between host and parasite richness"

**Supplemental Materials**

*Appendix A. Supplemental Methods*

The study employed a randomized complete block design, consisting of five spatial blocks, each 15 × 15 m (225 $m^2$). In each block, we established 64 plots, each 1 × 1 m with 1 m aisles between plots. In May 2011, the existing vegetation was removed from each plot using glyphosate herbicide (Riverdale® Razor® Pro, Nufarm Americas Inc, Burr Ridge, IL), but we did not apply herbicide to aisles between plots. Two weeks after herbicide application, we removed dead vegetation and covered all plots with landscape fabric. Each plot was then assigned to a combination of factorial treatments: native plant (i.e., host) richness with multiple native community compositions at each level of richness and soil nutrient supply.

*Host composition and species richness*

Each plot was assigned to one of two levels of initial host species richness: monoculture or five-species polyculture. These treatments represent the lower end of the natural range of variation in this system (Heckman et al 2016, 2017). Host species were selected from a pool of six native herbaceous perennials already present at Widener Farm. We selected host species that were present locally to ensure site suitability and to increase the likelihood that parasites capable of exploiting them were present locally. Our species pool included three grasses—*Andropogon virginicus, Setaria parviflora, Tridens flavus*, and three forbs—*Packera anonyma, Scutellaria integrifolia, Solidago pinetorum*.

Plants were propagated from seed in the greenhouse at the University of North Carolina at Chapel Hill then seedlings were transplanted into the soil through a small hole in the landscape fabric covering the plot. Each plot contained 41 individual plants, spaced approximately 10 cm apart in a checkerboard pattern. This planting density is low relative to the surrounding plant community. However, plant density recovers quickly in this system. By 2014, we observed comparable light attenuation between our experimental communities and unmanipulated parts of the same field, indicating that plant density had reached that of the surrounding vegetation (Heckman et al 2017). Polycultures contained 9 individuals of one randomly chosen species and 8 individuals of the other 4 species. In early summer 2012, we replaced all individual plants that had not survived the winter. *Setaria parviflora* was planted in 2012 instead of 2011, replacing a species that failed to establish in any plot in 2011. In July 2012, we removed landscape fabric from all plots, and removed non-planted individuals by hand.

In southeastern US old fields, succession toward woody dominance proceeds rapidly (Fridley and Wright 2012, Heckman et al. 2016). However, some disturbances, such as mowing and grazing, can arrest succession at the stage of dominance by herbaceous perennial species. In the North Carolina Piedmont, this successional stage is typical of two- or three-year old fields (Oosting 1942). In this study, we planted native perennial species that are common at this stage of herbaceous dominance (e.g., $C_4$ grasses and non-leguminous forbs) before allowing colonization to proceed. Because of the successional nature of the system, the pool of colonizers also changed over time. Specifically, early in the study, annual species that are typical of newly abandoned fields (e.g., *Gamochaeta purpurea*, *Conyza canadensis*) colonized the plots; these species were replaced by perennial species typical of later stages of old field succession, which were more common locally. Thus, over time, the species colonizing plots began to more closely resemble the surrounding community, leading to increasing similarity between the experimental plots and the surrounding vegetation (Heckman et al 2017).

*Quantification of final host community structure*

To avoid plot-level edge effects, we quantified the absolute cover of each species in a marked 0.75 × 0.75 m subplot in the center of each plot. Estimates of exotic plant abundance excluded several rare species (amounting to less than 1% of total cover in any plot) which were not identifiable to species.

The phylogeny of non-tree species, constructed using 'phyloGenerator' (Pearse & Purvis 2013), included options -gene *rbcL*, *matK* –alignment mafft –phylogen RAxML – integrated Bootstrap 1000, and constraint tree topology following Smith (2011). Sequence data were not available for *Asclepias syriaca* (which amounted to less than 1% of total cover in any plot), so the phyloGenerator function THOROUGH was used to replace sequences with the most closely related taxon using NCBI taxonomy. Plant phylogenetic diversity was calculated using the ses.mpd function in R package Picante (Kembel *et al.* 2010). Specifically, it was quantified using a null-modeling approach that measures the degree to which a plot is more or less phylogenetically diverse than random, given the number of host species and weighted by their relative abundance. To do this, we generated a z-score comparing the mean-pairwise-phylogenetic-distance between taxa in a plot to a randomly assembled plot with the same number and relative abundance of host species, permuted 1000 times. This allowed us to generate an estimate of host phylogenetic diversity that is independent of host species richness.

*Quantification of parasite richness*

Surveys of parasite richness of visually inspecting the five oldest leaves per host individual for damage by foliar parasites. On each leaf, parasites were categorized into morphospecies based on symptom morphology and fruiting body structures when visible (e.g.,

Liu *et al.* 2016; Halliday *et al.* 2017). Parasite morphospecies vouchers were either verified by the North Carolina State University plant disease clinic or by culturing fungal isolates from surface-sterilized lesions in 2% malt-extract agar. The cultured isolates were sorted using morphological characteristics, then DNA was extracted from each unique culture using a RED-Extract-N-Amp Plant kit (Sigma-Aldrich, St. Louis, Missouri, USA). The ITS region was amplified using the fungal-specific primers ITS 1F and ITS 4. The sequences were compared with those from GenBank using the UW-BLAST program (BLAST, http://blast.ncbi.nlm.nih.gov), yielding species names for a subset of fungal morphospecies (Table S1).

*Appendix B. Supplemental Results*

In our full longitudinal model of parasite richness, there was a positive relationship between host and parasite richness (post-assembly host richness $p=0.002$), this positive relationship depended on initial host diversity (initial host diversity × post-assembly host richness $p=0.04$), and this effect of initial host diversity changed over time (initial host diversity × post-assembly host richness × year $p=0.02$; Table S1). Fertilization reduced parasite richness by 10% (fertilization $p<0.001$), but this effect did not vary over time (fertilization × year $p=0.38$), did not affect the relationship between host and parasite richness (fertilization × post-assembly host richness $p=0.52$; fertilization × post-assembly host richness × year $p=0.81$), and did not alter the effect of initial host diversity on the relationship between host and parasite richness over time (initial host diversity × fertilization × post-assembly host richness × year $p=0.32$; Table S2). These non-significant interactions were therefore removed from the model, yielding a reduced model (Table S3).

Table S1. Parasite morphospecies and their associated host species grouped into four categories: A) planted host species, B) native colonizing host species, C) exotic colonizing host species, and D) host species with unknown geographic provenance. Parasite morphospecies is presented in the leftmost column, with parasite type in brackets and genbank accession numbers in parentheses.

**A) Planted host species**

| | *Andropogon virginicus* | *Setaria parviflora* | *Tridens flavus* | *Scutellaria integrifolia* | *Packera anonyma* | *Solidago pinetorum* |
|---|---|---|---|---|---|---|
| Balansia sp. [leaf stroma] | ■ | | | | | |
| Unidentified microbe [black leaf spot] | | | | ■ | | ■ |
| Phyllachora graminis [tar spot] | ■ | | | | | |
| Unidentified fungus [choke] | ■ | | | | | |
| Colletotrichum cereale [anthracnose] (MG016017, MG016018, MG461206, MG461207) | | | ■ | | | |
| Drechslera sp. [leaf spot] (MG016015, MG461208, MG461209) | ■ | | | | ■ | |
| Unidentified galling insect 1 | | | | | ■ | |
| Unidentified galling insect 3 | | | | | | ■ |
| Unidentified leaf mining insect | | | | | ■ | |
| Mycosphaerellaceae [leaf spot] | | ■ | ■ | | | |
| Rhizoctonia solani [leaf spot] | | ■ | | | | |
| Unidentified microbe [scorch] | | | | ■ | | ■ |
| Drechslera / Aureobasidium / Gaueumannomyces complex [leaf spot] (MG016008, MG016019, MG016021) | | | ■ | | | |
| Drechslera / Curvularia complex [leaf spot] (MG016006, MG016007) | ■ | | | | | |
| Stagonospora sp. [leaf spot] | | ■ | ■ | | | |
| Unidentified tent caterpillar | | | | | ■ | |

**B) Native colonizing host species**

| | *Conyza canadensis* | *Dichanthelium dichotomum* | *Erigeron annuus* | *Eragrostis capillaris* | *Juncus anthelatus* | *Oxalis dillenii* | *Schizachyrium scoparium* | *Solanum carolinense* | *Symphyotrichum pilosum* |
|---|---|---|---|---|---|---|---|---|---|
| Alternaria alternata [leaf spot] (MG461213) | | | | ■ | | | | | |
| Unidentified microbe [black leaf spot] | ■ | | | | | | | | |
| Unidentified fungus [leaf spot] | | ■ | | | | | | | |
| Phyllachora graminis [tar spot] | | | | | ■ | | ■ | | |
| Colletotrichum cereale [anthracnose] (MG016017, MG016018, MG461206, MG461207) | | ■ | | | | | | | |
| Unidentified fungus [leaf spot] | | | | | | | | | ■ |
| Drechslera sp. [leaf spot] (MG016015, MG461208, MG461209) | | | | | | | ■ | ■ | |
| Unidentified galling insect 3 | | | | | | | | | ■ |
| Unidentified microbe [scorch] | ■ | | | | | | | | |
| Unidentified tent caterpillar | | | | | | | | ■ | |

**C) Exotic colonizing host species**

| | *Anthoxanthum odoratum* | *Holcus lanatus* | *Lespedeza cuneata* | *Leucanthemum vulgaris* | *Lonicera japonica* | *Paspalum notatum* | *Plantago lanceolata* | *Rumex acetosella* | *Lolium arundinaceum* | *Sorghum halepense* |
|---|---|---|---|---|---|---|---|---|---|---|
| Bipolaris drechsleri [leaf spot] (MG461214) | | ■ | | | | | | | | |
| Unidentified fungus [leaf spot] | | ■ | | | | | | | | |
| Phyllachora graminis [tar spot] | | | | | | ■ | | | | |
| Colletotrichum cereale [anthracnose] (MG016017, MG016018, MG461206, MG461207) | ■ | | | | | | | | ■ | |
| Pestalotiopsis / Diaporthe complex [leaf spot] (MG461216, MG461219) | | | | | ■ | | | | | |
| Didymella glomerata [leaf spot] (MG461217, MG461218) | | | ■ | | | | | | | |
| Drechslera sp. [leaf spot] (MG016015, MG461208, MG461209) | ■ | | | | | | | | ■ | |
| Colletotrichum sublineolum / Alternaria / Aureobasidium complex [leaf spot] (MG461210, MG461211, MG461212) | | | | | | | | | | ■ |
| Puccinia coronata [rust] | | ■ | | | | | | | ■ | |
| Puccinia graminis [rust] | | | | | | | | | ■ | |
| Rhizoctonia solani [leaf spot] | ■ | ■ | | | | | | | ■ | |
| Sclerotinia sp. [white rot] (MG016009) | | ■ | | | | | | | | |
| Unidentified tent caterpillar | | | ■ | | | | | | | |

**D) Host species with unknown geographic provenance**

| | *Unknown Poaceae 2* | *Carex sp.* | *Dichanthelium sp. 2* |
|---|---|---|---|
| Unidentified microbe [chlorosis] | | | ■ |
| Unidentified fungus [yellow rust] | | ■ | |

**Table S2.** ANOVA for full longitudinal mixed model.

| | DF | F-value | p-value |
|---|---|---|---|
| Intercept | 1, 210 | 1911.03 | <0.0001 |
| Block | 4, 102 | 1.02 | 0.40 |
| Initial host diversity | 1, 10 | 49.27 | <0.0001 |
| Fertilization | 1, 102 | 18.81 | <0.0001 |
| Host species richness | 1, 210 | 10.18 | 0.0016 |
| Year | 2, 210 | 7.39 | 0.0008 |
| Initial diversity × Resources | 1, 102 | 9.42 | 0.0027 |
| Initial diversity × Host species richness | 1, 210 | 4.22 | 0.041 |
| Resources × Host species richness | 1, 210 | 0.42 | 0.52 |
| Initial diversity × Year | 2, 210 | 49.62 | <0.0001 |
| Resources × Year | 2, 210 | 0.97 | 0.38 |
| Host species richness × Year | 2, 210 | 2.35 | 0.10 |
| Initial diversity × Resources × Host species richness | 1, 210 | 3.13 | 0.078 |
| Initial diversity × Resources × Year | 2, 210 | 1.39 | 0.25 |
| Initial diversity × Host species richness × Year | 2, 210 | 3.92 | 0.021 |
| Resources × Host species richness × Year | 2, 210 | 0.21 | 0.81 |
| Initial diversity × Resources × Host species richness × Year | 2, 210 | 1.16 | 0.32 |

**Table S3.** ANOVA for reduced longitudinal mixed model.

| | DF | F-value | p-value |
|---|---|---|---|
| Intercept | 1, 220 | 2213.68 | <0.001 |
| Block | 4, 102 | 1.007 | 0.41 |
| Initial host diversity | 1, 10 | 57.06 | <0.0001 |
| Fertilization | 1, 102 | 17.97 | <0.0001 |
| Host species richness | 1, 220 | 9.94 | 0.0018 |
| Year | 2, 220 | 7.10 | 0.0010 |
| Initial diversity × Resources | 1, 102 | 9.31 | 0.0029 |
| Initial diversity × Host species richness | 1, 220 | 4.16 | 0.043 |
| Initial diversity × Year | 2, 220 | 49.08 | <0.0001 |
| Host species richness × Year | 2, 220 | 0.88 | 0.42 |
| Initial diversity × Host species richness × Year | 2, 220 | 4.71 | 0.010 |

**Table S4.** Coefficient estimates from the full structural equation model. Estimates are provided both raw and standardized to a common scale to facilitate comparisons. Correlations among dependent variables are indicated by ~~.

| Response | Predictor | Estimate | Std Error | z-value | p | Std Estimate |
|---|---|---|---|---|---|---|
| Rarefied parasite richness | Host Species Richness | 0.119 | 0.035 | 3.411 | 0.001 | 0.304 |
| | Host Exotic Abundance | 0.180 | 0.126 | 1.430 | 0.153 | 0.156 |
| | Host Phylogenetic Diversity | -0.293 | 0.131 | -2.234 | 0.025 | -0.193 |
| | Richness × Exotic Abundance | 0.082 | 0.021 | 3.842 | 0.000 | 0.255 |
| | Richness × Phylogenetic Diversity | 0.031 | 0.037 | 0.844 | 0.399 | 0.064 |
| Host Species Richness | Diversity | 1.341 | 0.877 | 1.529 | 0.126 | 0.197 |
| | Resources | -3.624 | 0.814 | -4.451 | 0.000 | -0.531 |
| | Diversity × Resources | -0.191 | 1.074 | -0.177 | 0.859 | -0.024 |
| Host Exotic Abundance | Diversity | -0.188 | 0.237 | -0.793 | 0.428 | -0.081 |
| | Resources | 1.402 | 0.311 | 4.503 | 0.000 | 0.603 |
| | Diversity × Resources | -0.522 | 0.373 | -1.400 | 0.162 | -0.197 |
| Host Phylogenetic Diversity | Diversity | 0.407 | 0.221 | 1.843 | 0.065 | 0.231 |
| | Resources | -0.211 | 0.275 | -0.768 | 0.443 | -0.120 |
| | Diversity × Resources | 0.199 | 0.319 | 0.624 | 0.533 | 0.099 |
| ~~ Host Species Richness | ~~ Host Exotic Abundance | -0.634 | 0.262 | -2.424 | 0.015 | -0.232 |
| ~~ Host Exotic Abundance | ~~ Host Phylogenetic Diversity | -0.158 | 0.088 | -1.789 | 0.074 | -0.192 |

Goodness of fit: Robust $X^2$ p=.29, RMSEA p=.618, SRMR=.075

**Table S5.** Coefficient estimates from the final structural equation model. Estimates are provided both raw and standardized to a common scale to facilitate comparisons. Correlations among dependent variables are indicated by ~~.

| Response | Predictor | Estimate | Std Error | z-value | p | Std Estimate |
|---|---|---|---|---|---|---|
| Rarefied parasite richness | Host Species Richness | 0.119 | 0.035 | 3.395 | 0.001 | 0.303 |
| | Host Exotic Abundance | 0.180 | 0.126 | 1.426 | 0.154 | 0.156 |
| | Host Phylogenetic Diversity | -0.293 | 0.131 | -2.232 | 0.026 | -0.193 |
| | Richness × Exotic Abundance | 0.082 | 0.022 | 3.788 | 0.000 | 0.255 |
| | Richness × Phylogenetic Diversity | 0.031 | 0.037 | 0.845 | 0.398 | 0.064 |
| Host Species Richness | Diversity | 1.244 | 0.532 | 2.338 | 0.019 | 0.182 |
| | Resources | -3.721 | 0.511 | -7.284 | 0.000 | -0.546 |
| Host Exotic Abundance | Diversity | -0.454 | 0.189 | -2.401 | 0.016 | -0.195 |
| | Resources | 1.136 | 0.188 | 6.052 | 0.000 | 0.489 |
| Host Phylogenetic Diversity | Diversity | 0.508 | 0.153 | 3.313 | 0.001 | 0.289 |
| | Resources | -0.110 | 0.157 | -0.701 | 0.484 | -0.063 |
| ~~ Host Species Richness | ~~ Host Exotic Abundance | -0.630 | 0.263 | -2.398 | 0.016 | -0.229 |
| ~~ Host Exotic Abundance | ~~ Host Phylogenetic Diversity | -0.164 | 0.089 | -1.844 | 0.065 | -0.198 |

Goodness of fit: Robust $X^2$ p=.222, RMSEA p=0.436, SRMR=.086

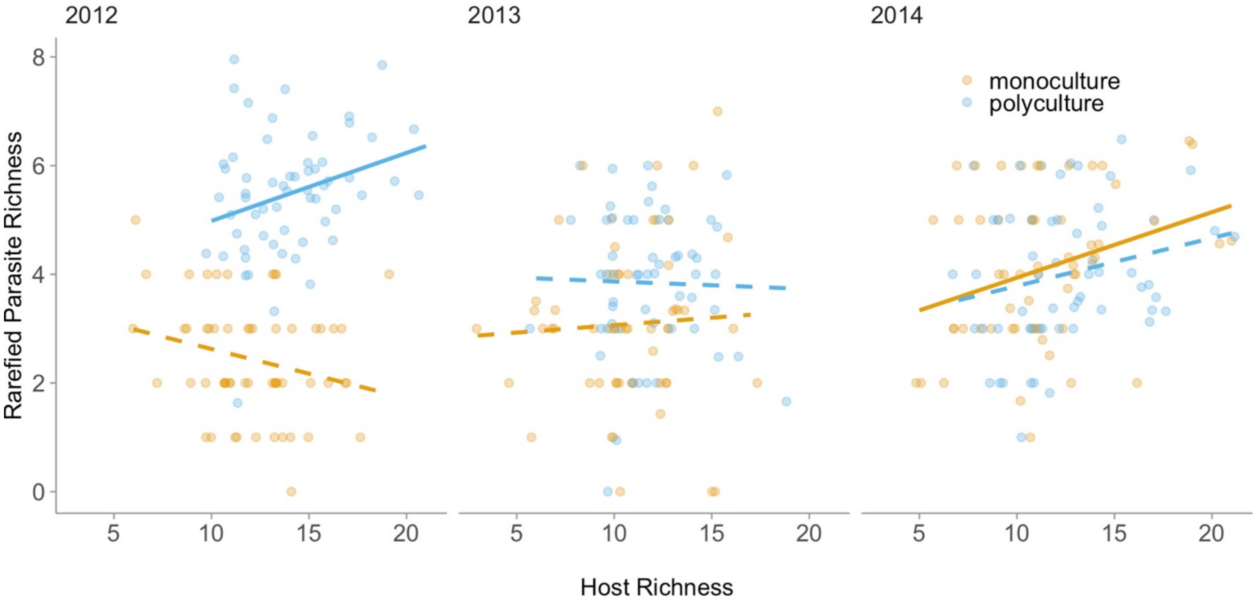


**Figure S1.** Longitudinal mixed model results showing the model-estimated effect of host richness on rarefied parasite richness as a function of initial host richness (monocultures orange; polycultures blue) across three years. The y-axis shows rarefied parasite species richness, and the x-axis shows contemporaneous host richness. Points are the raw data, and lines represent the model-estimated effect of the experimental treatment on the relationship between host and parasite-richness (i.e., the interactive effect of contemporaneous host richness and initial host diversity on parasite richness). Solid lines are significantly positive ($p<0.05$), while dashed lines are not significantly positive ($p>0.05$). Caution should be taken when comparing model estimates among years as sampling methodology differed among years. The positive relationship between host and parasite richness that was observed in polyculture plots in 2012 weakened over time, while the relationship between host and parasite richness in monoculture plots strengthened over time, resulting in a positive relationship in 2014.

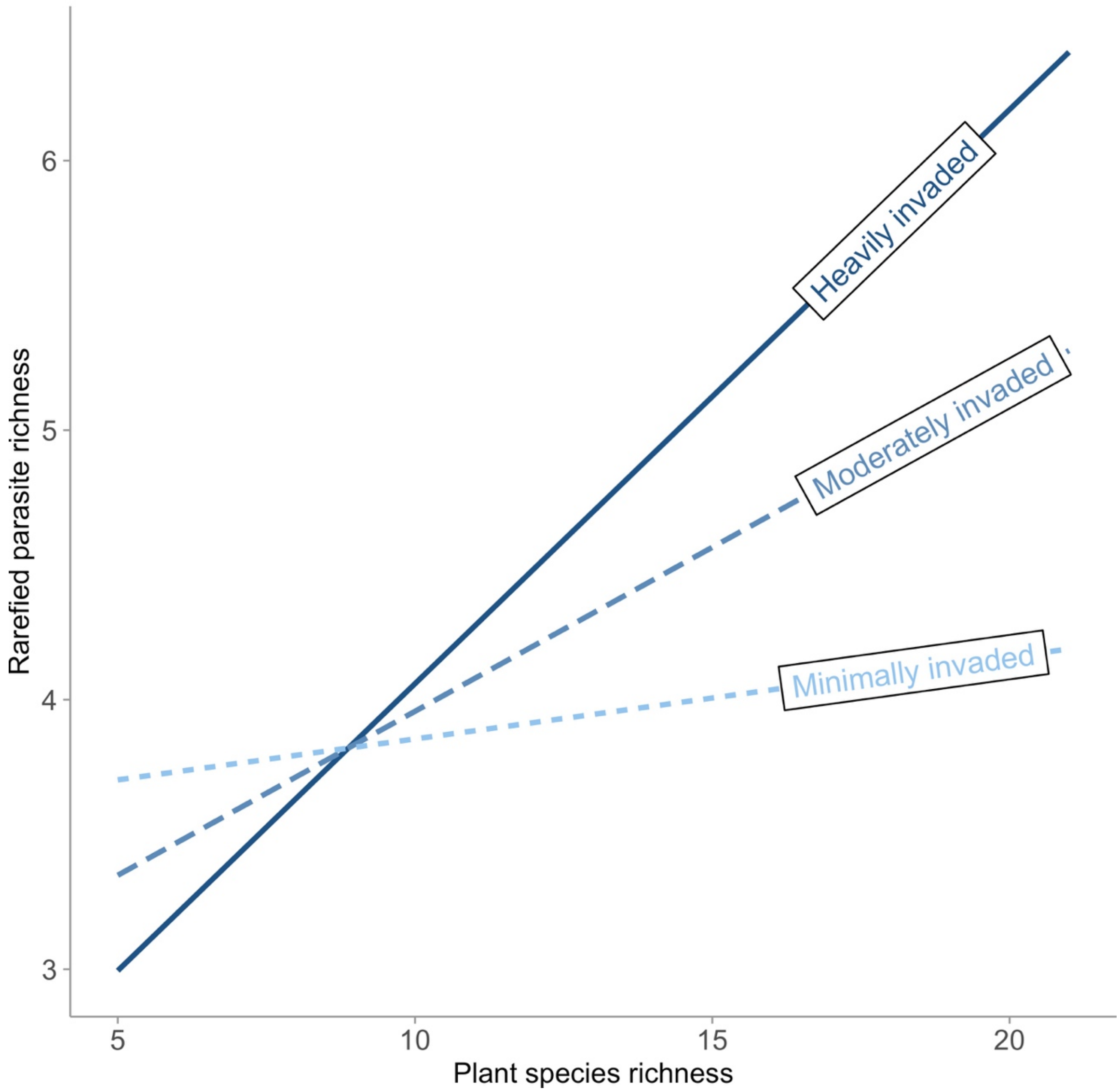


**Figure S2.** Simple slopes analysis showing the effect of host richness on parasite richness as a function of exotic host abundance. Model estimated effects of standardized and centered exotic abundance on the slope of the relationship between final host richness and parasite richness (i.e., the interactive effect of final host richness and exotic host abundance on parasite richness). “Heavily invaded” communities are one standard deviation greater than the mean of exotic abundance. “Moderately invaded” communities are at the mean of exotic abundance across the experiment. “Uninvaded” communities are one standard deviation below the mean of exotic abundance. Communities that became most dominated by exotic species also exhibited the strongest positive relationship between host and parasite richness.
